# Telemonitoring System Architecture for Emotion Recognition with WBSN and Ensemble Learning

**DOI:** 10.1101/2022.07.25.501385

**Authors:** Maryam El Azhari

## Abstract

Telemonitoring provides a set of technologies that enables remote monitoring of patients with critical health conditions.Wireless Body Area Networks (WBAN) technology has emerged as a major factor contributor to the need for ubiquitous healthcare.It provides a real-time telemonitoring system to treat several chronic diseases using wireless biosensors.The first standard to support communication between biosensors is IEEE 802.15 Task Groups 6 (TG6). The standard regulates the communication in,on or inside the human body for a short-range and low power wireless communications.IEEE 802.15.6 operates on a Medium Access Control (MAC) layer that supports several Physical (PHY) layers such as narrowband (NB),ultra-wideband (UWB),and human body communication (HBC).In this paper,a realtime telemonitoring system architecture for EEG brainwave emotion recognition is presented.The system aims to collect EEG data and forward them to clinicians in order to detect the evolution of the monitored diseases based on brain wave emotion analysis.Ensemble Learning techniques are used for decision making to discern the current health state of the patient.

## 1 Introduction

Wireless Body Sensor Networks is a subform of telehealth that provides a remote patient monitoring [1].This technology allows patients to perform daily activities while a stream of physiological data measurements is continuously communicated to clinicians in medical centers. [2]. A WBSN consists of several biosensors capable of measuring numerous medical parameters such as: EEG signal, ECG signal, body temperature and arterial pressure.The measured parameters are transmitted to the base station where medical analysis are performed.The overall architecture of a WBSN is made of three level architecture. the first level includes the communication from biosensors to the Personal Digital Assistant (PDA) and vice-versa. The PDA is usually located within the transmission range of all biosensors. The second level consists of a wireless technology that connects the PDA to the internet.It can be either a WIFI, a 4G or a Wireless Sensor Network [3]. The third level includes a set of unconstrained-power devices linked to a distant mega-database where patients medical records are stored for analysis [4]. Although WBSNs prove their promising impact on providing a full coverage of patients heath status throughout the day, there are still many design requirements to be taken into account. Examples of such design requirements are:

- **Communication range**:it should be enough to allow sensors around the body to communicate with each other, a minimum of 2-meter communication range should be feasible.
- **Low power**:biosensors should operate on a low power to prevent unwanted heating of the skin. This problem can lead to severe health complications.To resolve this issue, biosensors should operate under regulatory set of rules to minimize the heating measured by the specific absorption rate (SAR) and keep it under a safe-threshold value.
- **Coexistence**:WBSNs should be endowed with adaptable mechanisms to coexist with other wireless technologies operating in the same spectrum like ZigBee, Wireless Sensor Networks, Bluetooth and Wireless Local Area Networks.
- **Interference**: WBSN is more susceptible to collision, mutual interference and external interference caused consecutively by the high mobility of the subject, coexistence with other neighbouring WBSNs and the existence of other technologies such as WiFi or Bluetooth.The interference contributes to a considerable inefficiency of the channel performance.

Although many wireless technologies have been developed to respond to the design requirements of WBSNs, there still exist many issues to tackle such as energy efficiency, varying data rates, dynamic topology, security and QoS. The medical data received at the base station i.e health care servers or data warehouses are further analyzed by specialist to provide medical decisions and intervene when necessary. The recent growth of Artificial Intelligence (AI) field has equipped many companies Worldwide with intelligent decision-making machines. A large area of AI is dedicated to Machine Learning(ML) which refers to the ability of a computer to learn from mined datasets.Ensemble Learning has been widely used in machine learning models.It combines multiple classifier systems for decision making. Ensemble systems have being proven to be uniquely beneficial and greatly flexible when dealing with real-world application problems. In this paper, a realtime telemonitoring system architecture for EEG brainwave emotion recognition is presented.The system aims to collect EEG data and forward them to medical servers for analysis purposes.Ensemble Learning methods are applied to detect the evolution of the monitored diseases based on the brain wave emotion analysis.

## 2 Overview

There are several mental illnesses that alter the mood and emotions in different ways. Examples of such diseases are: schizophrenia, depression, insomnia, attention deficit hyperactivity disorder(adhd), sleep disorder, epilepsy and autism. With the rapid progress of technologies in neural control interface (nci),also referred to as braincomputer interface (bci), it has become possible to capture the brainwave signals and proceed with emotion classification using machine learning algorithms.

### 2.1 The importance of electroencephalogram in emotion detection

Electroencephalography (EEG) records electrical activities of neural cells, it helps to detect distinct states from scalp surface area. The signal frequencies are generally ranging from 0.1 Hz to more than 100 Hz and they are categorized into alpha,beta,theta,delta,gamma and theta signals. Pattern recognition of EEG signal enables to detect the emotional psychology of an individual in real time.The area of the brain that is involved in controlling emotions, learning and memory is referred to as the limbic system (Figure 2).

**Fig. 1:**
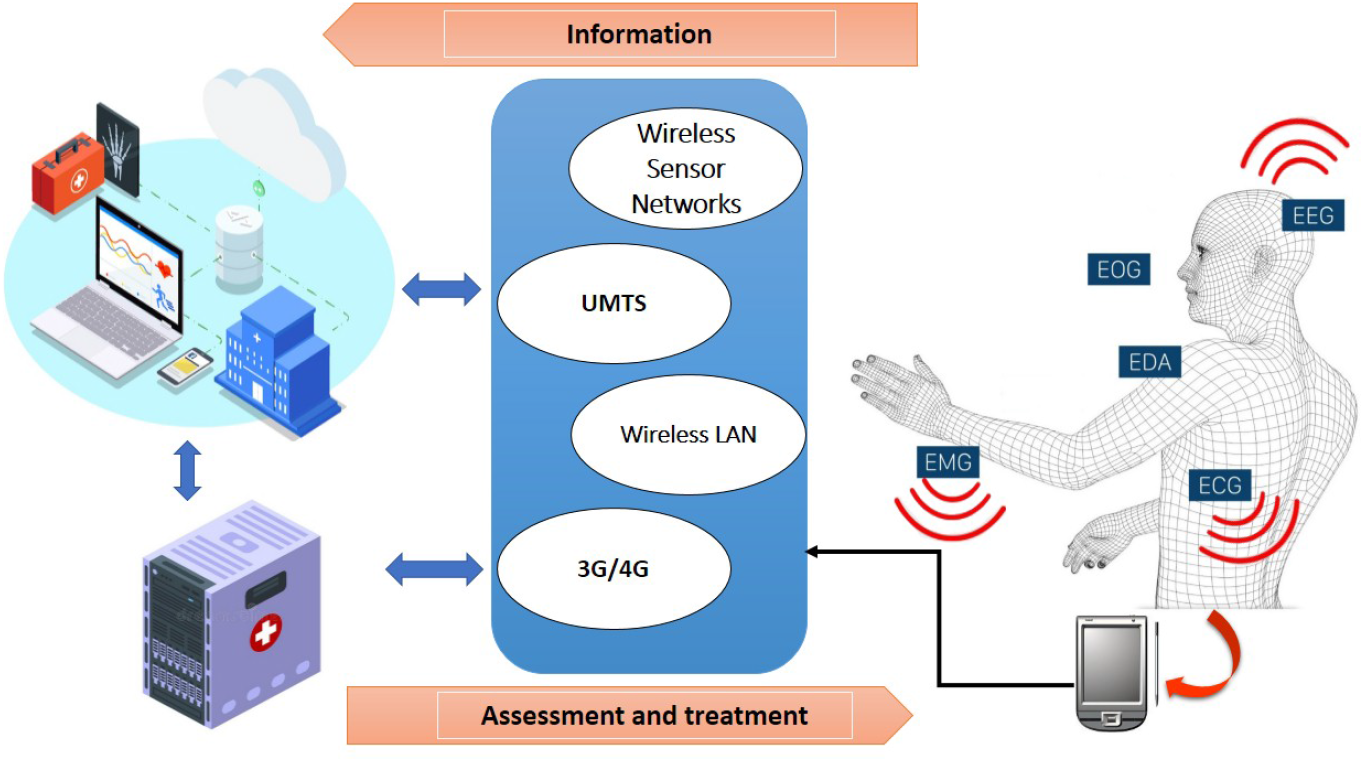
Three level architecture of a WBSN [5]

**Fig. 2:**
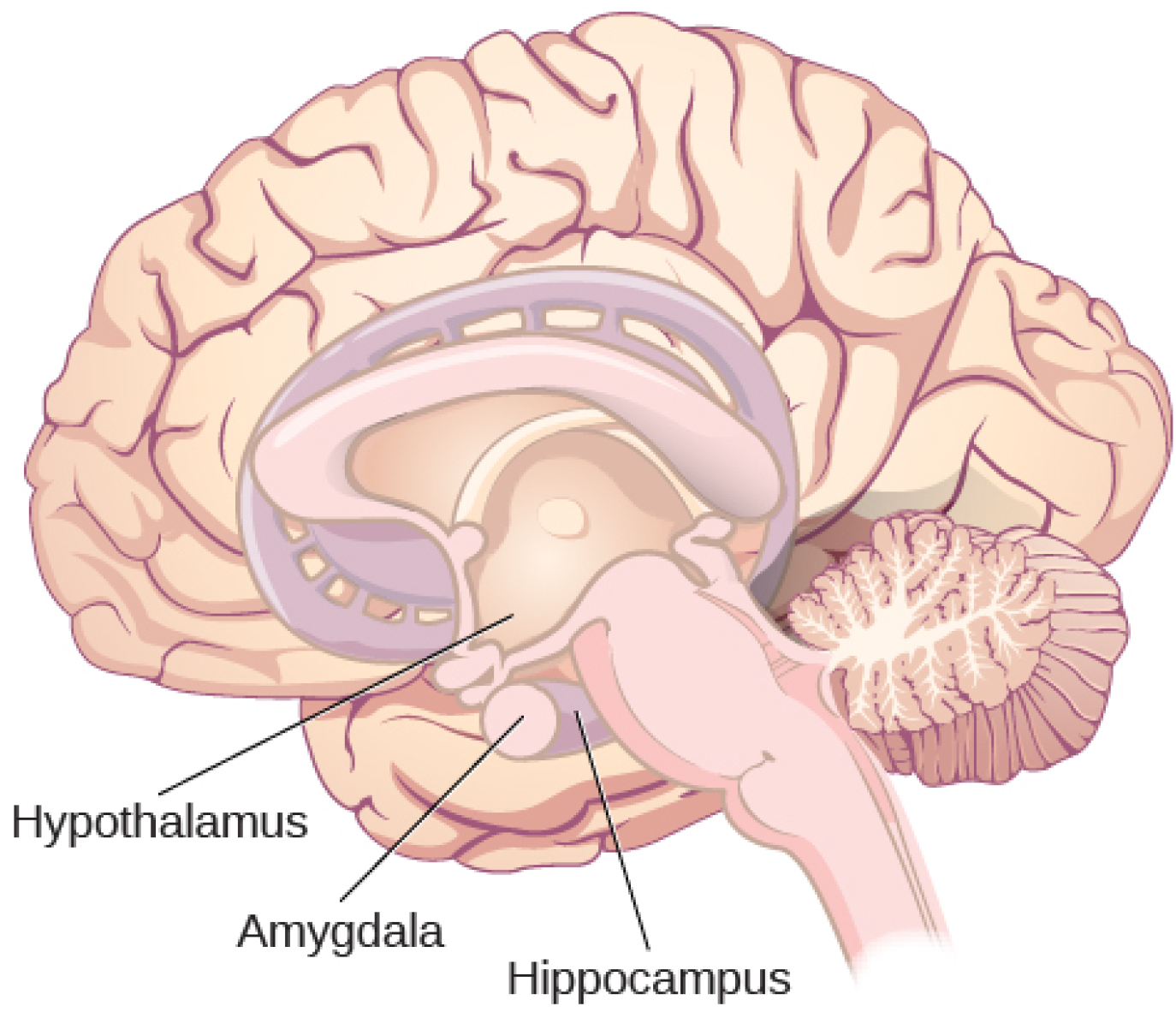
The limbic system includes the hypothalamus, amygdala, and the hippocampus, and is involved in generating emotional response and memory [6]

The limbic system incorporates the hypothalamus, amygdala, hippocampus and thalamus. The hypothalamus plays a part in generating emotional reaction by activating the sympathetic nervous system. The amygdata contributes to the emotional information processing and its transmission over cortical structures.The thalamus is considered a sensory relay center whose primary function is to relay sensory signals to the cerebral cortex.Hippocampus is an important component of the limbic system, it regulates emotion, learning, and memory.Hippocampus is very sensitive and it is one of the first areas to be affected by mental disorders such as depression, anxiety, schizophrenia and phobias. The International Federation of Societies for Electroencephalography and Clinical Neurophysiology (IFSECN) has given many standard rules on electroencephalography in clinical practice.IFSECN recommended the conventional 10–20 system [7] where the 21 electrodes are distributed uniformly(Figure 3.a),thus covering the human scalp.A higher-resolution systems include extra electrodes to record a more detailed electroencephalography. These electrodes cover intermediate sites between the existing 10–20 system (Figure 3.b) [8].

**Fig. 3:**
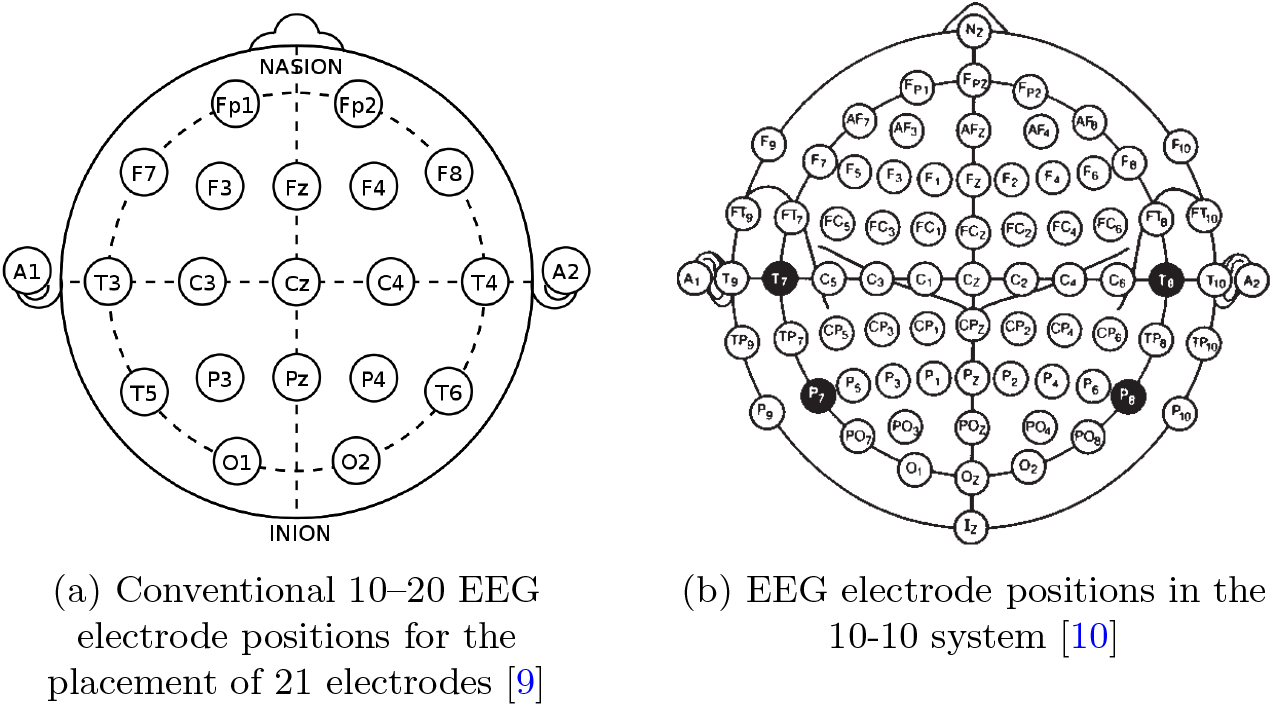
EEG electrode positions in 10-x system

### 2.2 Wireless EEG biosensors

The advancement in wireless communication and electronics manufacturing has enabled the development of wireless EEG biosensors. These EEG biosensors allow patients to exercise daily activities while being under medical supervision. A common trait of wireless biosensors is the constrained energy usage.Wireless biosensors operate on a low powered batteries, therefore it is very crucial to use optimized energy-based communications paradigms to extend the biosensor lifetime. Researchers from university of Illinois have created a new class of electronic sensors in the form of biomedical implants to monitor patients after a brain injury or surgery.These brain biosensors have the advantage of self-dissolving when they are no longer required, thus mitigating the risk of infection due to monitors removal [11].Yu M. Chi and al. utilized non-contact electrodes (Figure 4) to record high quality EEG/ECG signals in [12].The proposed wireless EEG/ECG system offers many advantages since non-contact electrodes can function without an ohmic connection to the body.The non-contact electrodes raise the limitations of wet and dry electrodes embodied in direct contact to the skin and discomfort of use.

**Fig. 4:**
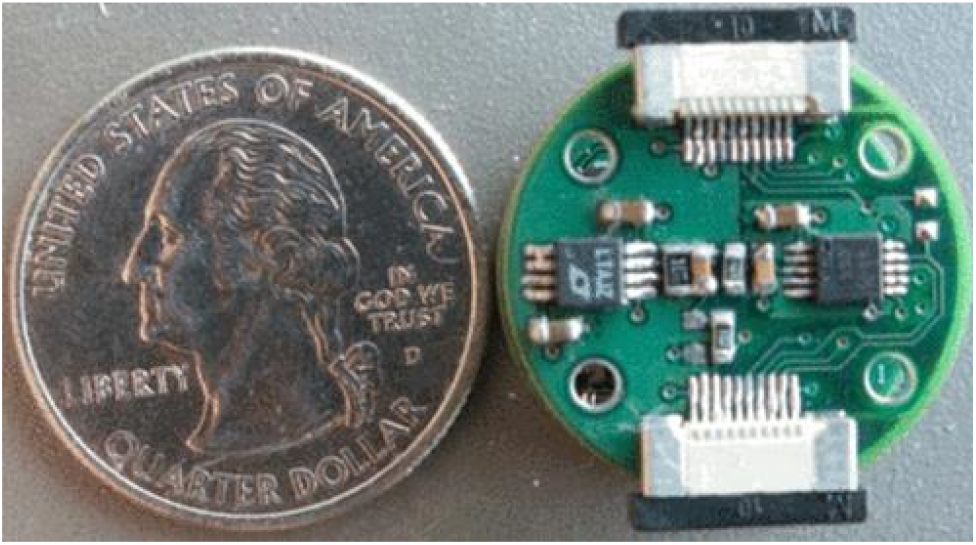
Non-contact electrodes [13]

### 2.3 IEEE 802.15.6

IEEE 802.15.6 is a standard presented by (TG6) to regulate communication in Wireless Body Sensor Networks. The standard aims to enhance the services of medical and health care systems. There are several physical layers on which IEEE 802.15.6 operates, such as human body communication, ultra-wideband, narrow band, etc. Two types of topologies can be used for IEEE 802.15.6 : single hop topology and multi-hop topology. With a single hop topology, the frames are sent directly to the hub, whilst the multi-hop topology includes a mediator set between the biosensors and the hub to exchange the frames. In this case study, the single hop topology is used for communication.The hub coordinates the access of multiple sending nodes to the channel, through three different access modes:

- Beacon mode with beacon period superframe boundaries.
- Non-beacon mode with superframe boundaries.
- Non-beacon mode without superframe boundaries.

A superframe is divided into multiple time slots, where each slot has a ranking number. An example of the superframe structure can be shown in Figure 5. The superframe may include several periods such as : Random Access Phase (RAP),Exclusive Access Period (EAP),Contention Access Period (CAP) and optionally a B2 frame. When the length of any phrase is set to null, then the phase is automatically eliminated. CSMA/CA and Slotted ALOHA are two examples of random channel access used for contention [14].

**Fig. 5:**
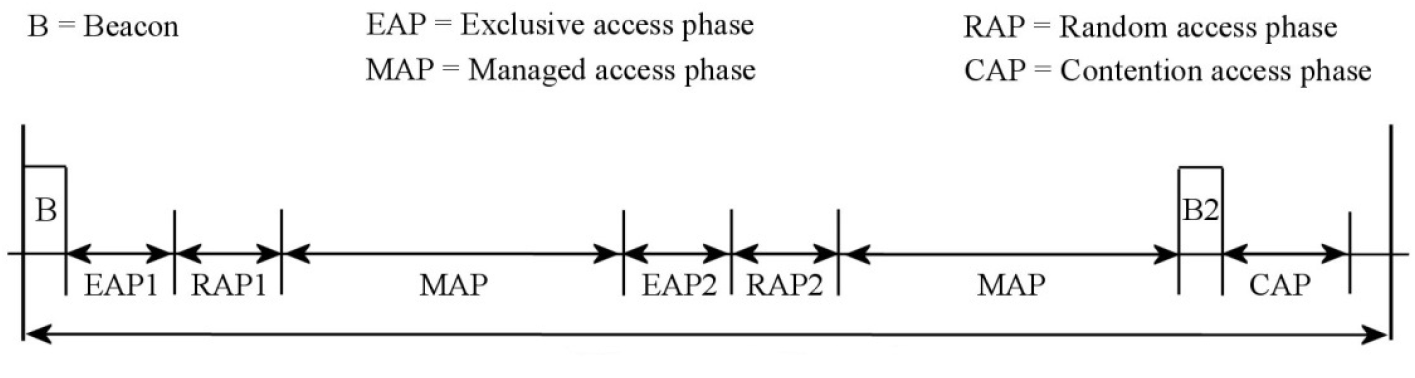
Superframe structure of 802.15.6 MAC [15]

In [16],H.Renwei and al. gave an analytical comparison of the IEEE 802.15.4 and 802.15.6 Wireless Standards based on MAC layer.The most significant features of both standards were highlighted while considering the different aspects of frequency, data rate and communication range. In [17], Z. Zhao and al. proposed a queuing analytical framework to evaluate the performance of IEEE 802.15.6. The utilized scheduling algorithm is CSMA/CA with some specific constraints such as : instantaneous delay and body shadowing effects. A buffer-overflow queuing paradigm was developed to approximate the over-deadline packet dropping by solving a buffer length optimization problem. The authors in [17] [18] investigated the MAC and PHY layers of WBAN nodes as described in IEEE 802.15.6. The access mechanisms and the different types of communications supported by the standard had also been described.The conducted study revealed that the data flow and data security remain challenging in heath-care related applications.The performance of IEEE 802.15.6 CSMA/CA MAC had been analyzed under non-ideal channel condition using DTMC model in [19].It is concluded that the highest priority node provides better reliability, high throughput, low latency, and lower CCA failure probability. In [20], the effect of user priorities (UPs) on the performance of IEEE 802.15.6 CSMA/CA channel access mechanism was evaluated in narrow band. Simulation metrics mainly focus on the normalized throughput and average packet delay in which the traffic arrival rate and traffic distribution vary. The simulation results demonstrated that the user priorities mechanism makes a big influence on the network performance, it guarantees reliable and timely service for higher UPs while sacrificing the performance of lower UPs. It was also concluded that employing the CSMA/CA scheme alone is not enough to meet all needs of medical traffic. To this end, other access modes in IEEE 802.15.6 such as the improvised access and scheduled access were to be considered for further analysis.

## 3 Dynamic-priority based schema for channel access

In this section, a new contention channel access schema based on a dynamic priority is presented.The schema consists in introducing a tolerable data reception threshold above which data reception rate is acceptable.The proposed schema is mainly destined to biosensors with normal and low-priority as emergent data requires instant access to the channel. The Dynamic-priority based schema inherits its functioning mechanism from the CSMA/CA which likewise regulates the access to the channel in accordance with the type of traffic.In order to get a new allocation, the biosensor uses a backoff counter and a contention window (CW) as depicted in equation (1).The CW_*i*_ is computed based on the number of successful data packets reception and the priority of the biosensor. A high value of *CW*_*i*_ leads to a high probability to access the channel and conversely. For each *biosensor*_*i*_, the backoff counter is initialized to a random integer from [0, CW_*i*_] (equation (3)). The biosensor considers a slot idle if the slot is inactive for pCCATime period.The backoff counter is decremented by *step*_*i*_ for every idle CSMA slot. *step*_*i*_ (see equation (2))takes into consideration the value of *CW*_*i*_ and the maximum value that a contention window could reach for both medical and non-medical services. When the backoff counter value is equal to zero, data packet (e.g. management packet) is transmitted by the biosensors. However, when the channel is detected to be busy, the backoff counter is locked by the biosensor until the end of the current packet transmission.Similarly, when the time is insufficient to finish the current transmission, the backoff counter is blocked. Contrastingly, the backoff counter is unlocked when the channel is idle for the pSIFS period within the CAP and RAP, and also when the time is sufficient to finish the current transmission. In order to ensure a high data reception rate,three thresholds are introduced: *Success_Max,Failure_Max* and *priority*_*min*_. As described in algorithm 1, the new schema monitors the number of successful data packet transmission *Success*_*i*_ and regularly compares it to *Success*_*Max*_ before accessing the channel. If *Success*_*i*_ overcomes the value of Success Max threshold,then the priority of *biosensor*_*i*_ is increased. In this study,the biosensor with high priority_*i*_ is considered to hold nonemergent traffic. A *priority*_*i*_ is decreased when the biosensor_*i*_ fails to transmit data or management packets for a certain number of times that overcomes *Failure_Max*.The frequent change of *priority*_*i*_ leads to an equal contention for the channel access.

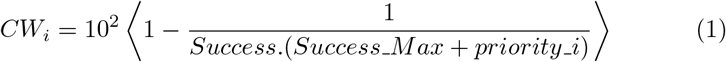

- *CW*_*i*_ : Contention Window of sensor node i.
- *Success_Max*: the maximum number of successful data packet reception.
- *Success* : the number of successful data packet reception.
- *priority*_*i*_ : the priority of sensor node i.

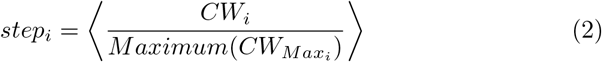

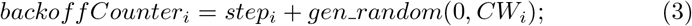
- *gen_random*(0, *a*) : a function that generates a random number between 0 and a

#### Algorithm 1

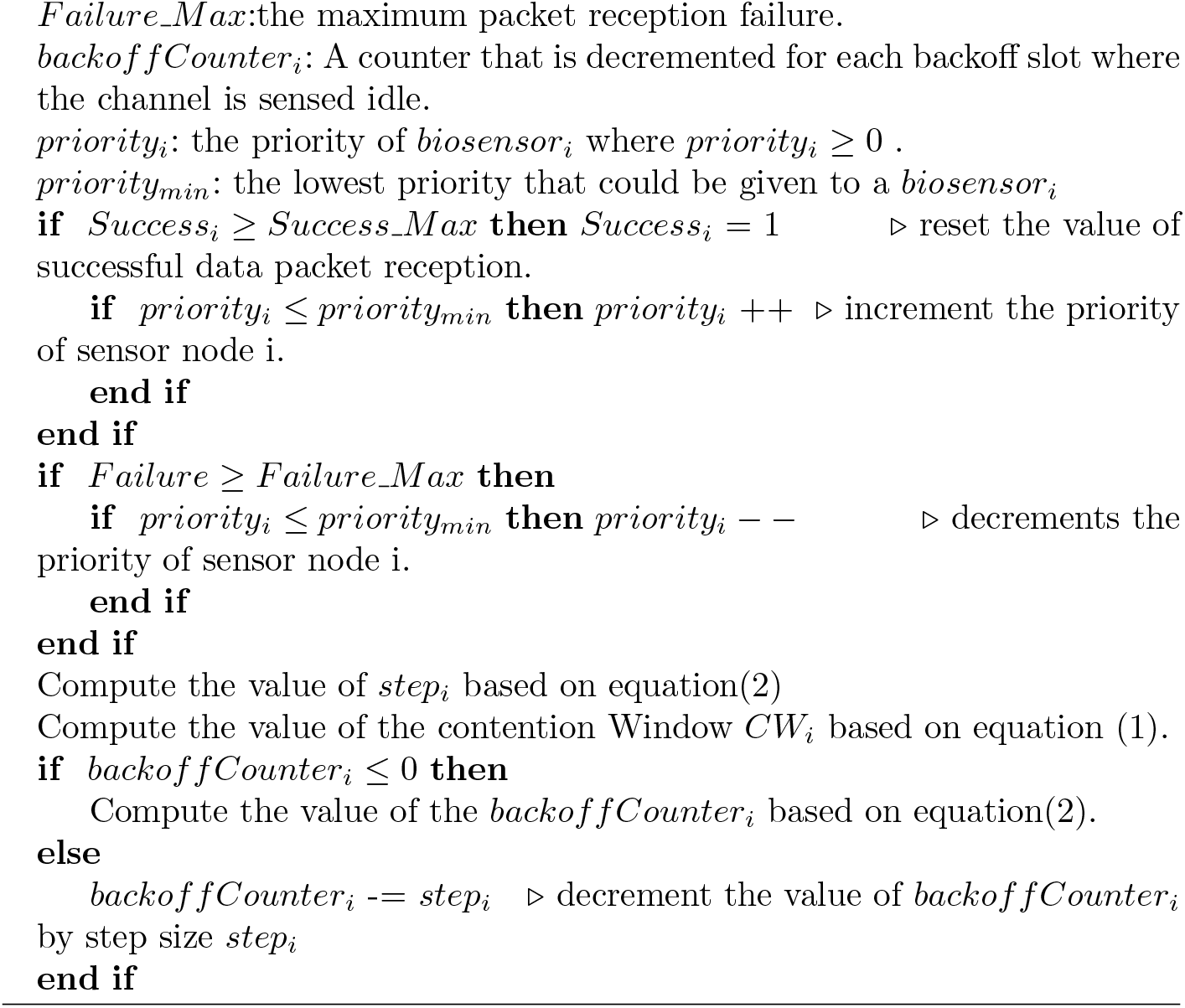

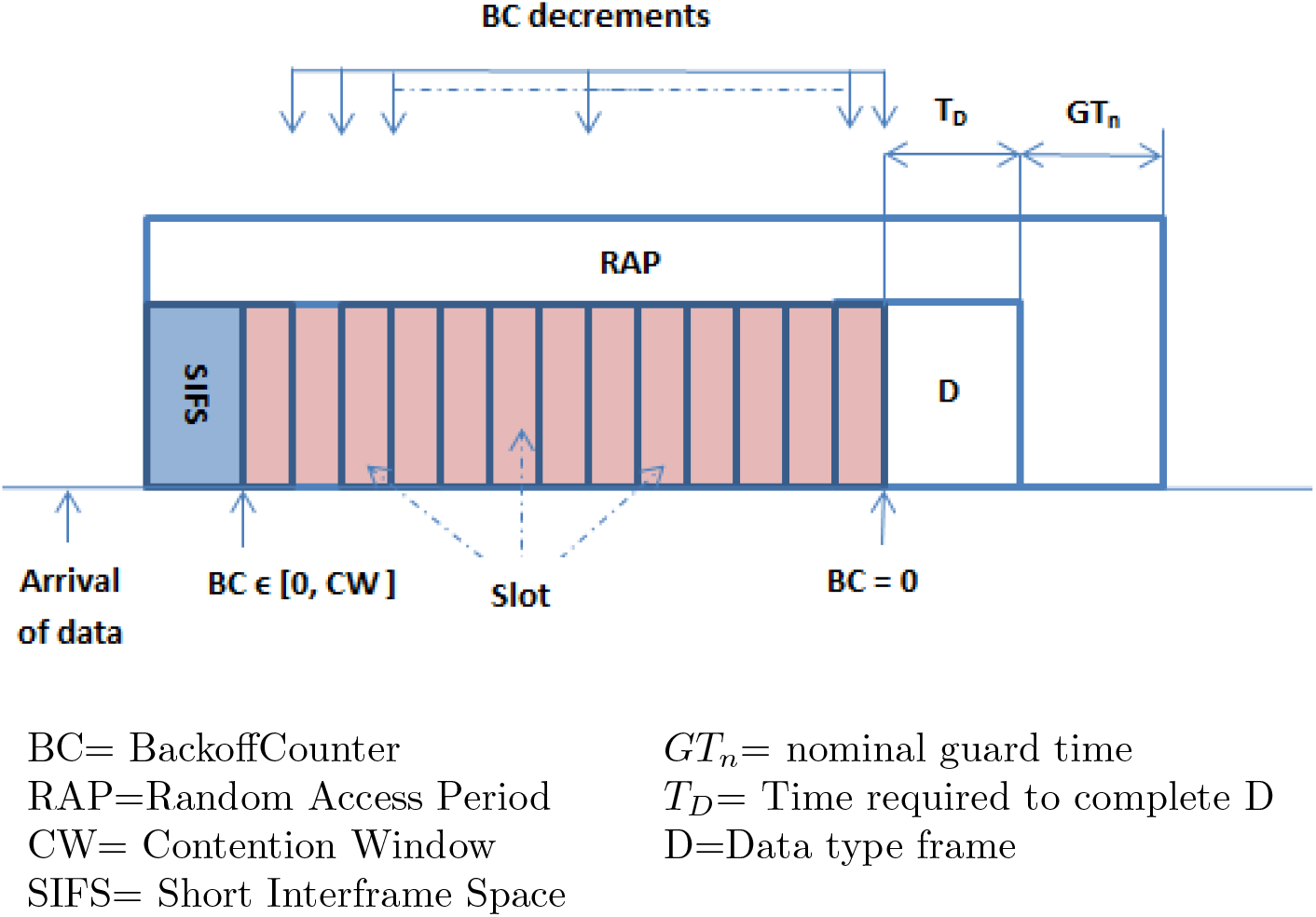

### 3.1 Simulation results

The simulation is performed in Castalia 3.3,which is an OMNET++ based simulation framework designed for WBSNs and WSNs.The efficiency of the proposed scheme is evaluated based on packet reception and end to end delay.The simulation parameters considered in this study are described in [21],where one coordinator node is located on the right hip, and five sensor nodes (at the extremities of the four limbs and the chest) regularly send packets of 130bytes (including overhead) to the coordinator.The average path loss between every pair is calculated from the experimental measurements [21].The packet generation rate is constant and equals to 5 pps (packets/sec) whereas the initial energy level of each sensor node is estimated to equal 18720J [22]. The average temporal variation of the channel and the path loss map are defined in Castalia distribution (pathLossMap.txt and TemporalModel.txt).The radio reception mode is set to “high”which is equivalent to 1024 for data rate (kbps), DIFFQPSK modulation type,bandwidth (MHz), sensitivity (dBm) and power consumed (mW) have respectively the following values: 20,-87 and 3.1. The beacon period of IEEE 802.15.6 is set to 32 slots where the first slot is dedicated to sending beacon packet for sensors synchronization and new connections management.The beacon slot is followed by the Random Access Period (RAP) period composed of 8 slots.During this period, biosensors contend to access the channel using a contention-resolving algorithm. In this case study,the rest of the superframe is set to null,thus allowing the transceivers to turn off their radio for for battery durability [21].As can be shown in Figure 6 and Figure 7,the new schema mitigates the average packet failure rate at the medium access control and radio modules respectively,which indicates a successful reception of data and management packets.Packet reception failure is caused by many factors from which we distinguish: interference,collision and non-reception state of the transceiver module.When the human body experiences a high mobility,the connection between the sender and the receiver is frequently corrupted,leading to bandwidth loss due to unused time slots.Moreover,an important number of packets are subject to intensive collision at the hub level(i.e. the PDA),and to interference between biosensors due to a false declaration of channel availability during the contention period.When the hub is carried by the human subject,it can be within or out of the transmission range of the biosensors,this results in enhancing the possibility of Human Body Shadowing,and causes a low sensitivity and a high packet loss rate.Another key factor that can be taken into account is the intrusion attack [23].An intruder may inject the networks with exponential number of fake packets to create collision,the intruder can also maintain a low priority to keep a sustainable control over the channel for a prolonged period of time.The duty cycle of all biosensors is identical for both schema as no synchronization schema is utilized,this explains the constant value of the average energy consumption among all biosensors that is equal to 0.15797 (nJ/bit). Figure 8 indicates the data packet latency at the application layer level. As can be shown in Figure 8,a large chunk of data is received in less than 20 ms for both static and dynamic priority-based schemes.It can also be observed that this amount decreases over time with preference given to a dynamic-based priority throughout several chunks of time intervals.This behavior was expected as the average packet reception with a dynamic-based priority scheme is relatively high.Moreover,the proposed scheme aims to guarantee an equal access to the channel while preserving the contention access mode,which results in a slight delay overhead.The non-negligible portion of packets at [600 .. int) bucket indicates an oncoming saturation. [24] [25].

**Fig. 6:**
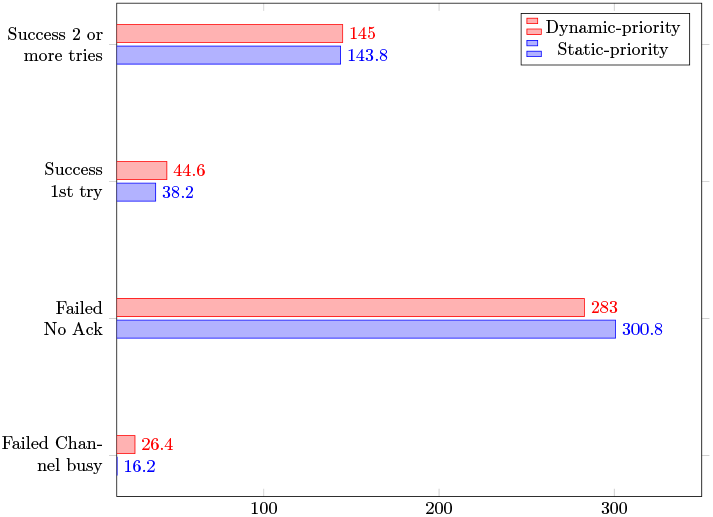
Data packet breakdown

**Fig. 7:**
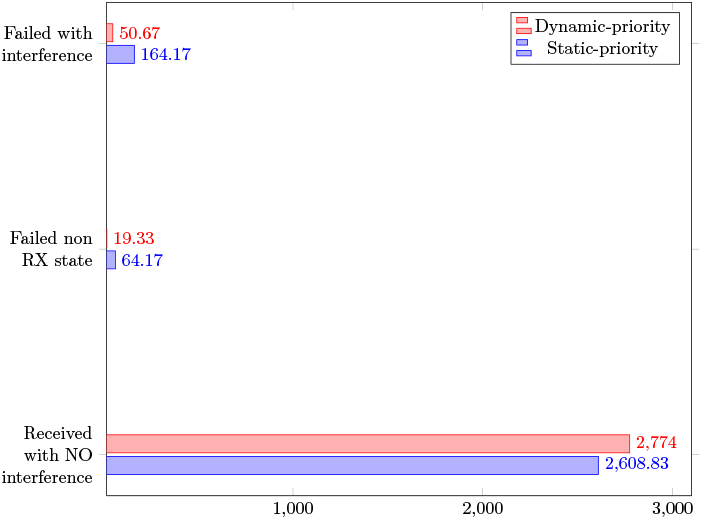
Rx packet breakdown

**Fig. 8:**
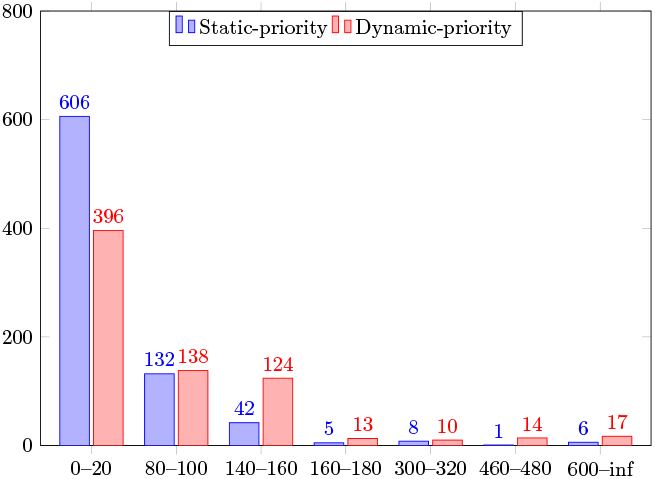
End-to-end latency

## 4 EEG data analysis

Upon receiving EEG data on healthcare data storage, analyses are further conducted to detect the evolution of the monitored diseases based on EEG emotion recognition.In this paper,ensemble learning is applied to perform EEG analyses owing to its advantage in achieving better predictive performance over a single model,but also reducing the dispersion of the predictions [26].

### 4.1 Ensemble learning

Ensemble learning is a combination of multiple machine learning models with the aim of providing an improved average performance over a single machine learning model. Building an effective ensemble learning system is based on three main pillars: data selection,training classifiers separately and combining member classifiers[27].Ensemble learning methods are divided into the following two categories: sequential ensemble methods and parallel ensemble methods.Sequential ensemble methods consist in generating a sequence of weak learners (or decision stumps) to exploit the dependence between weak learners.In contrast, parallel ensemble methods combine a set of base learners generated in parallel in order to exploit the independence between the base learners. There are three main types of ensemble learning methods:bagging,boosting and stacking.

- **Bagging**: this ensemble technique is based on bootstrap sampling to generate different base learners. Bootstrapping divides the dataset into subsets of n samples,where each subset constitutes the training data for a weak machine learning model.The sampling of each subset is done with replacement techniques to ensure randomization of the selected samples.Aggregation in bagging joins the results of training individual weak machine leanings models into a single decision.The methods used to make a final decision is voting in case of classification and calculating the average when dealing with regression (Figure 9).
- **Boosting**:this ensemble technique is considered as the most effective extension of the decision trees. It is designed to provide a strong learner based on a sequence of weak classifiers with the aim of creating a near-perfect classifier.Boosting technique initially starts with training a weak learner on a particular data subset.When the base learner .i.e decision stump shows a weak performance by getting some examples wrong,a new weak learner is embedded in the process to focus more on the weak points of the previous decision stump and resolve them. This process is repeated until a certain predefined condition is met and a final single strong learner is delivered at the end of boosting (Figure 10).
- **Stacking**: Stacking consists in using heterogeneous machine learning models to make predictions on the same dataset. Stacking defines two level learners: the first-level learners refer to individual learners that train the original dataset to generate a new training data set delivered to the second-level learner called meta-learner.This technique can also be thought of as stacking one layer of meta learner on top of another layer of base learners (Figure 11).

**Fig. 9:**
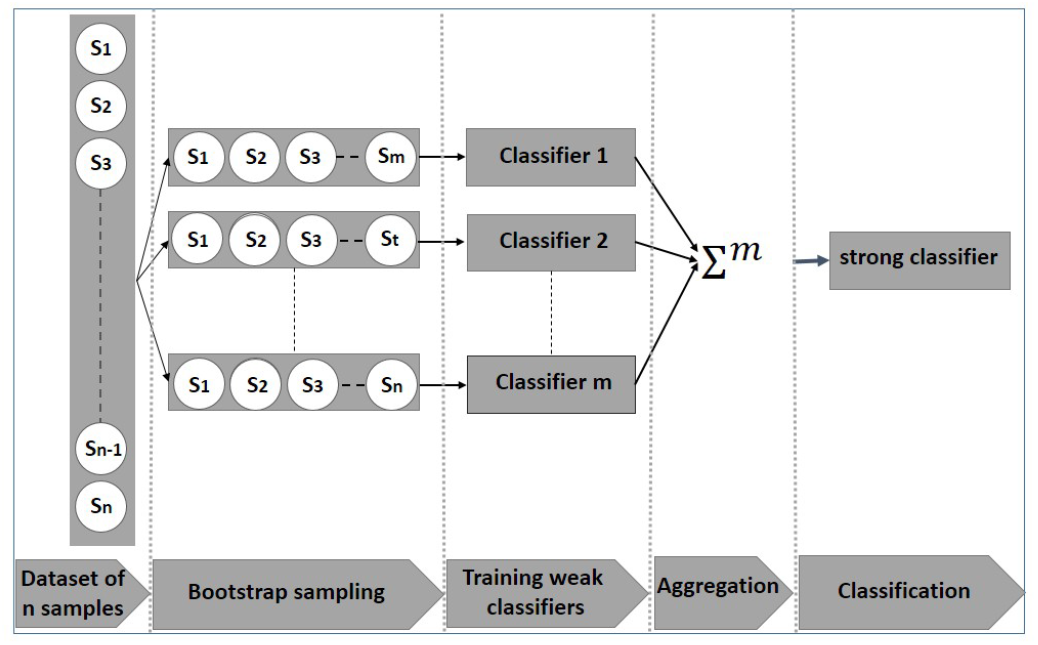
Bagging ensemble method

**Fig. 10:**
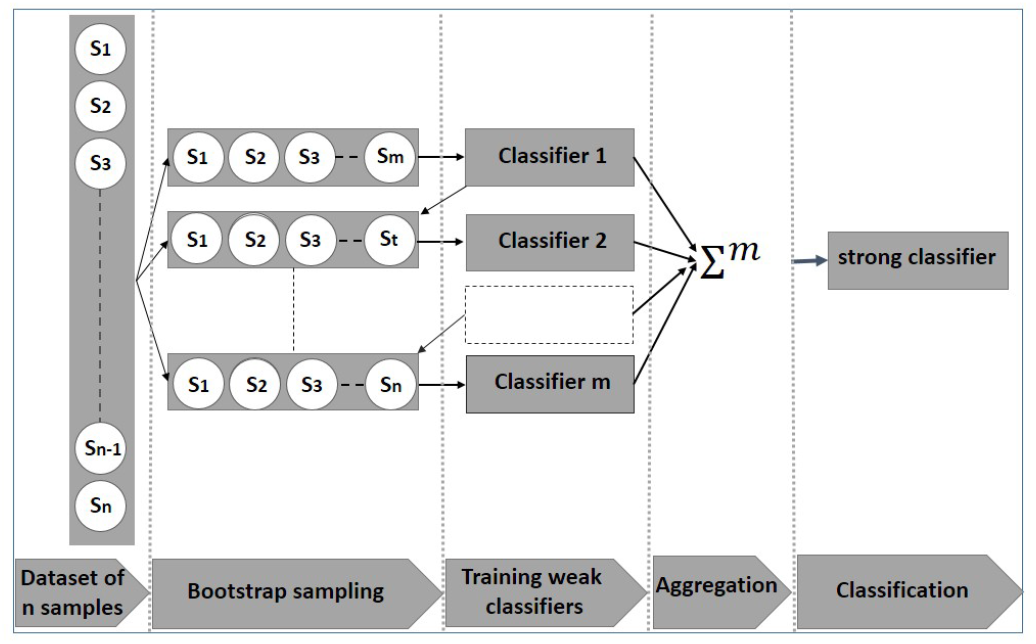
Boosting ensemble method

**Fig. 11:**
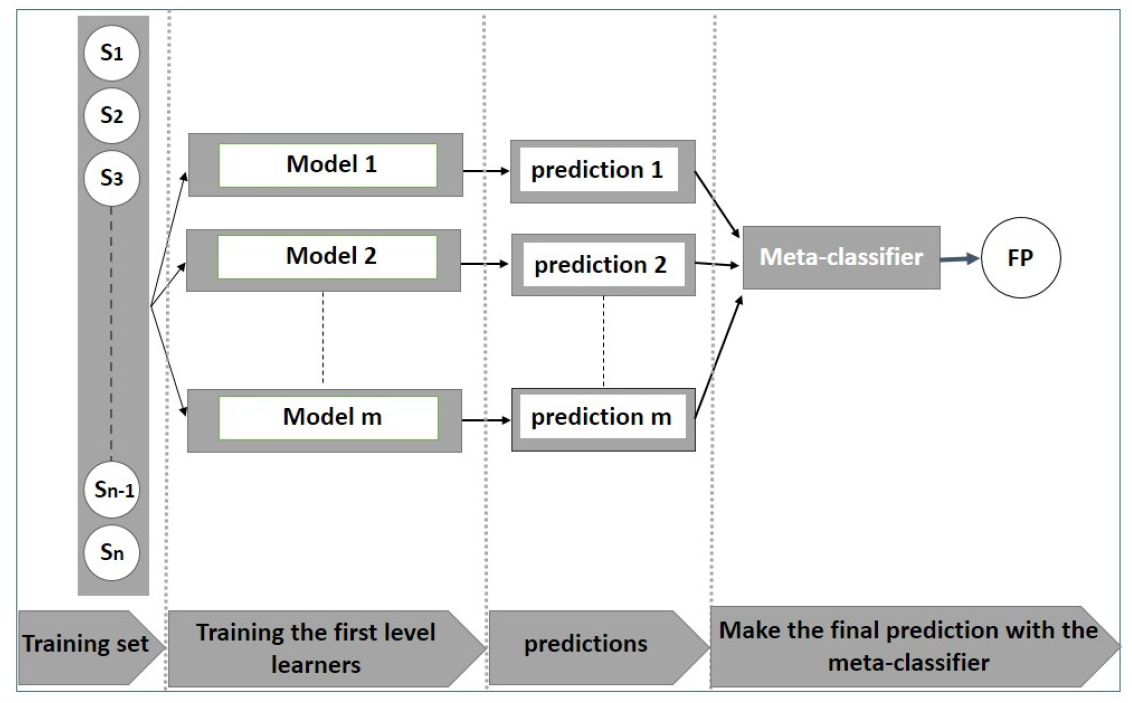
Stacking ensemble method

## 5 Experiments and results

The proposed real-time monitoring system starts with capturing a new trigger that either corresponds to a periodic or event-based data transmission (Figure 12).Upon receiving a new trigger,*α,γ,δ* and *θ* brainwaves are collected to be transmitted during Type-I/II phase of IEEE 802.15.6 MAC super-frame.Emergency traffic (which is also referred to as Type I data) is transmitted during the exclusive access phase (EAP-I/II) whilst non-emergency data (Type II data) are managed with random access phase (RAP-I/II) and contention access phase(CAP).EEG data is collected by the coordinator(i.e. the PDA) to be transmitted to the medical servers for analysis purposes,via an intermediate network such as: WSN,UMTS,WLAN,3G or 4G.

**Fig. 12:**
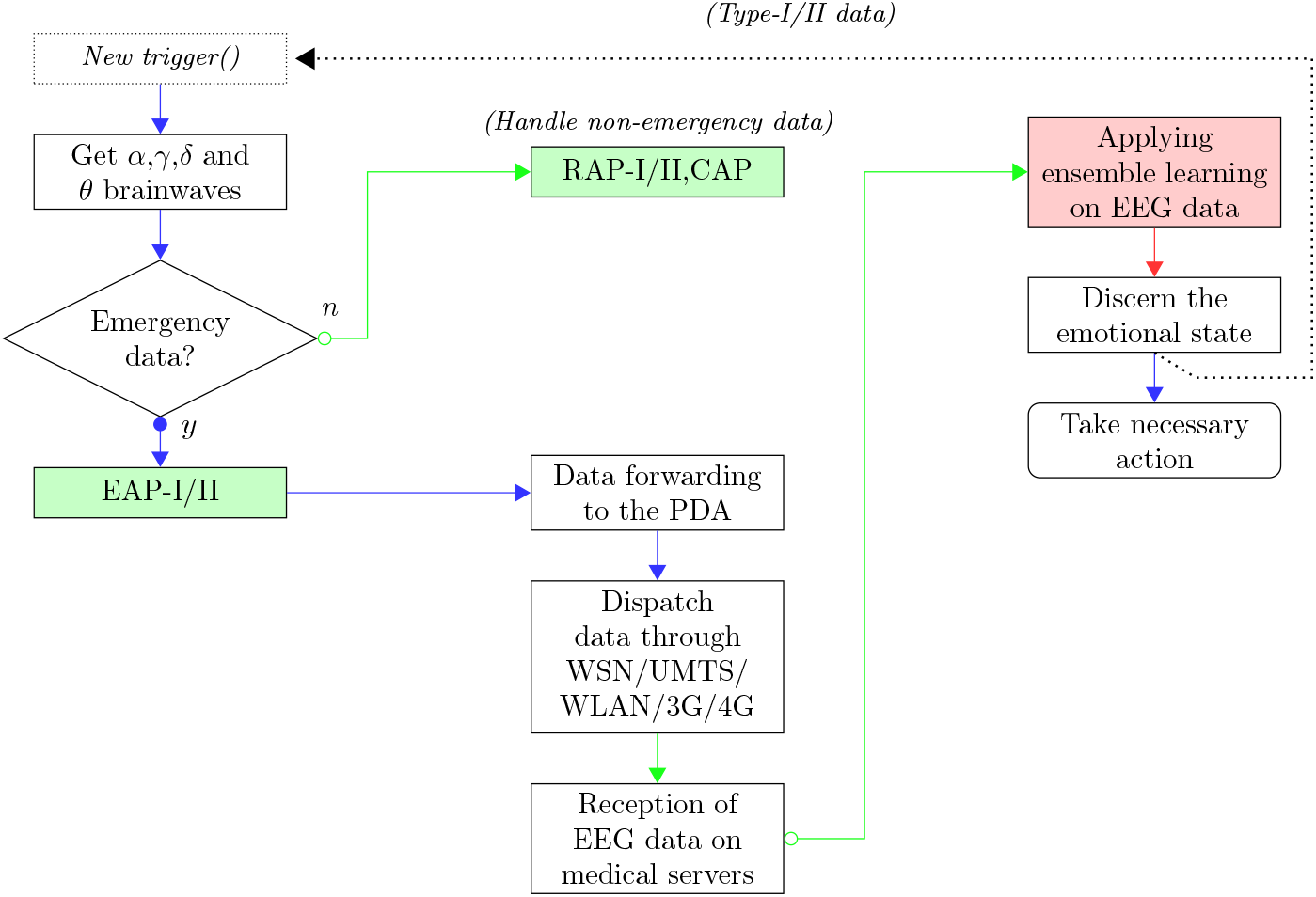
The organizational chart of the real-Time telemonitoring System

The study conducted in this paper uses the EEG brainwave dataset presented in [28] [29] to create an ensemble machine learning algorithm for the mental emotional state classification.The study presented in [28] employed four dry-electrodes embedded in a MUSE EEG headband.

Microvoltage measurements are taken from the TP9, AF7, AF8, and TP10 electrodes.The data were recorded from a 20 yo male and a 22 yo female for a time period T equals to 60s. Neutral data were collected without stimuli whilst positive and negative emotions were invoked using film clips. A total of 36 minutes of EEG data were recorded from the two subjects (6 minutes for each emotional state).Statistical extraction was further applied to generate a large dataset file that includes alpha, beta, theta, delta and gamma brainwaves. The organizational chart depicted in Figure 14 presents the machine learning algorithms selected to proceed with EEG brainwave classification.The three ensemble learning methods were considered in this study.The dataset is split into train and test sets with 80-20 ratio.The imbalance ratio of the dataset is 132:147:148,which is assumed to be slightly imbalanced and annul the use of oversampling and undersampling.The experiments are performed on a Virtual machine having 13GB RAM,33GB HDD and 2-core Xeon 2.2GHz.There are various evaluation metrics to assess the performance of the ensemble learning methods.All the following metrics used in this study are based on the confusion matrix:precision,recall,f1-score,accuracy,macro-average and weighted-average.The experimental results are summarized in Table 1. As can be discerned from the results,boosting ensemble methods slightly outperform bagging and stacking methods,however,the overall performance of the three ensemble methods is considered good as precision,recall and f1-score demonstrate a value greater than 90%. Hyperparameter tuning methods can also be combined with ensemble methods to find the best sets of predictive values,Random Search,grid search,bayesian optimization and tree-structured Parzen estimators are examples of such hyperparameter tuning methods.

**Fig. 13:**
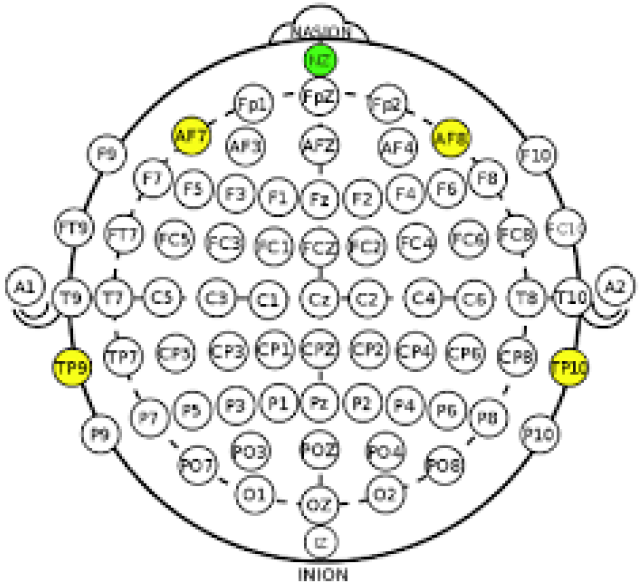
A MUSE EEG headband with TP9,AF7,AF8 and TP10 electrodes

**Fig. 14:**
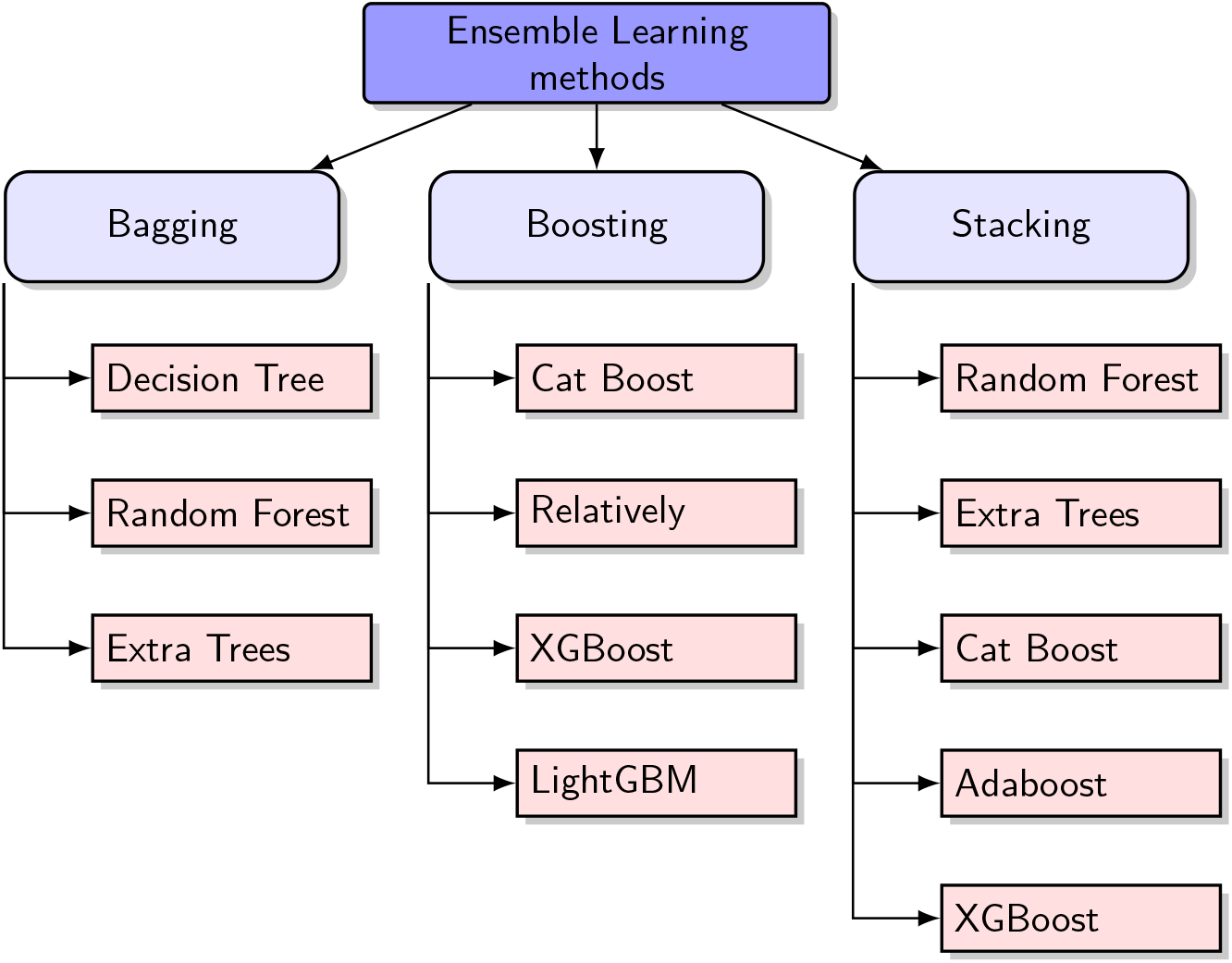
Ensemble machine learning organigram

**Table 1:**
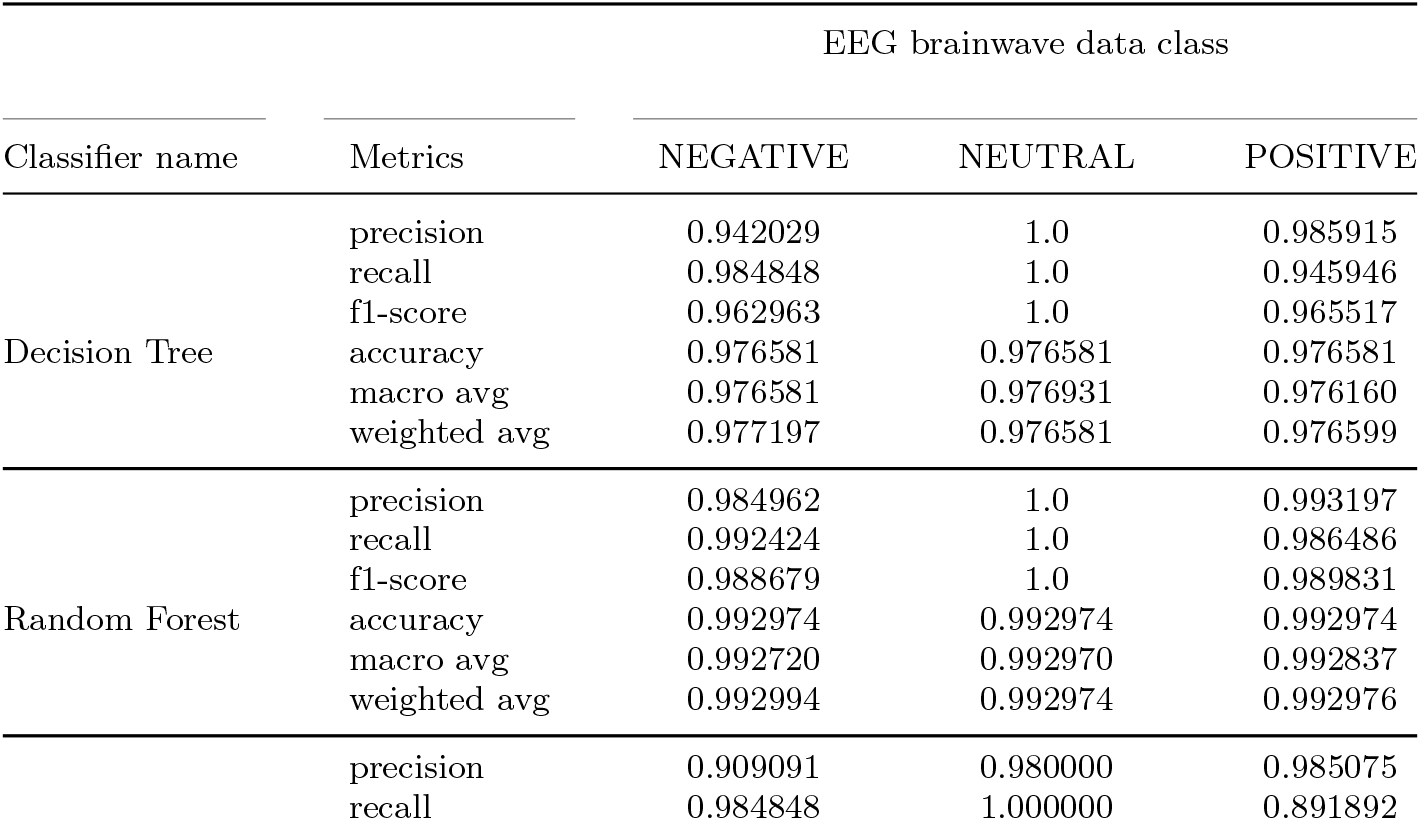
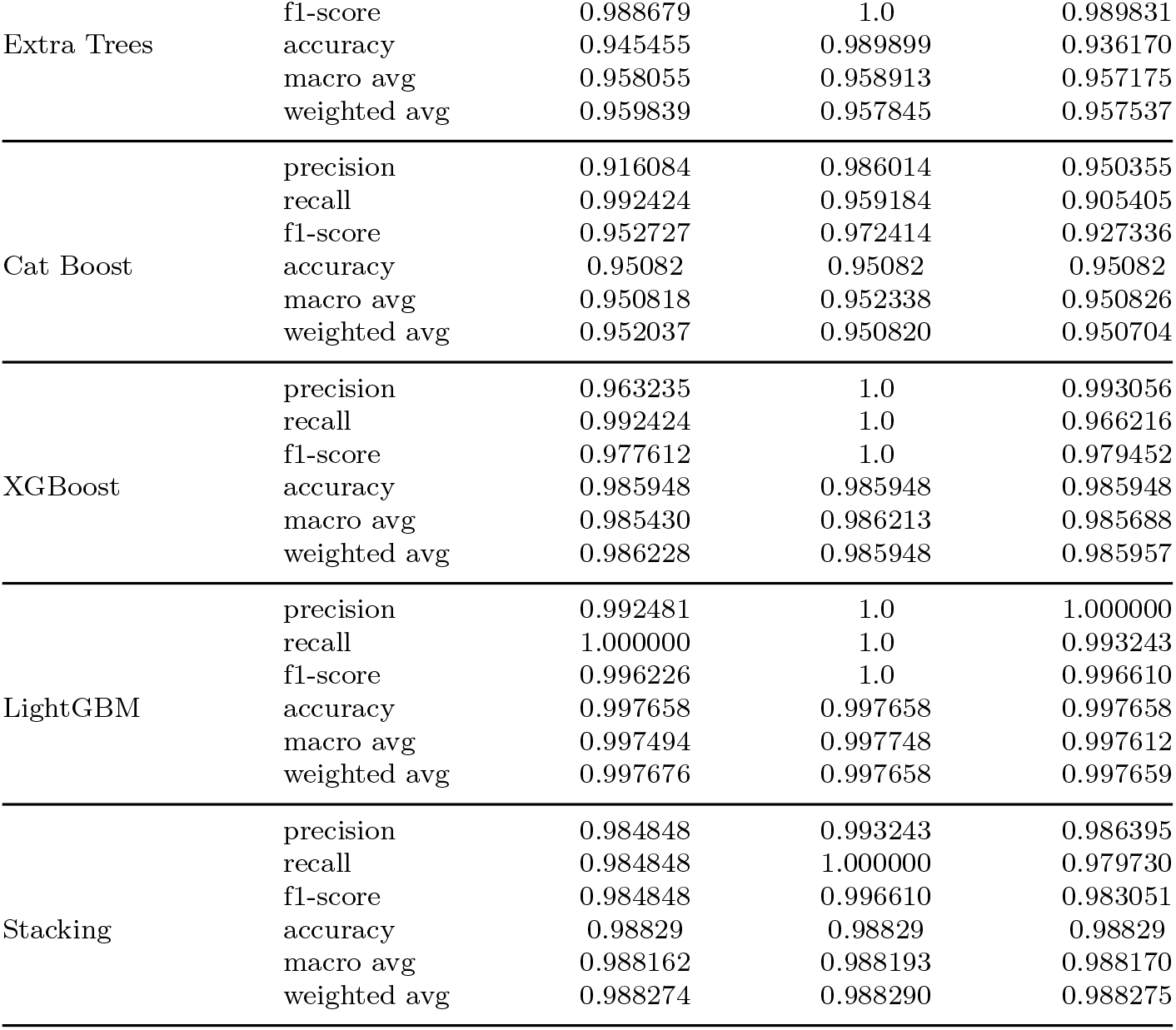
Experimental results of ensemble learning methods application on EEG brainwave dataset

## 6 Conclusion

A realtime telemonitoring system architecture is presented in this paper for EEG brainwave emotion recognition using Wireless Body Sensor Networks and ensemble learning. WBAN consists in using biosensors to measure physiological parameters and forward them to the base station for analysis purposes.A dynamic-priority based scheme for channel access regulation was proposed.The scheme defines new thresholds to guarantee an optimal use of the channel and increase the packet reception rate with respect to the traffic priority type.The simulation results have proven the efficiency of the proposed scheme in terms of mitigating packet failure due to collision,low sensitivity and busy state of the transceiver. A new classifier for EEG brainwaves detection will be created in future work to handle more complex EEG emotions for accurate disease diagnosis. An extended version of the proposed dynamic-priority scheme will also be studied in more detail to regulate the contention in high mobile environment,where multiple Wireless Body Sensor Networks coexist. The preliminary study highlights different challenges to be taken into account,including the management of the limited bandwidth for a perfect synchronization of biosensors,minimizing the overall energy consumption and finally reducing the end-to-end delay for both emergency and non-emergency traffic with a common channel access scheme shared among various WBSNs.

### Compliance with Ethical Standards

- Conflict of interest:Author Maryam El Azhari declares that he/she has no conflict of interest.
- Financial interests:The author has no relevant financial or non-financial interests to disclose.
- Ethical approval: This article does not contain any studies with human participants performed by any of the authors.

## References

[1] Yang, G.-Z.: Body sensor networks. Springer, London 72(2), 564 (2014)

[2] Yang, G.-Z.: Body Sensor Networks. Springer, Secaucus, NJ, USA (2006)

[3] Fahmy, H.M.A.: Wireless sensor networks. Springer, Singapore 1, 614 (2016). https://doi.org/10.1007/978-981-10-0412-4

[4] Khattiya, W., Timakul, S., Choomchuay, S.: An error control coding in mac layer for uwb wban. In: 2013 IEEE International Conference on Signal Processing, Communication and Computing (ICSPCC 2013), pp. 1–5 (2013). https://doi.org/10.1109/ICSPCC.2013.6664010

[5] https://biosignalsplux.com/products/sensors.html

[6] https://courses.lumenlearning.com/wmopen-psychology/chapter/reading-the-limbic-system-and-other-brain-areas/

[7] William O. Tatum, D. IV: Handbook of eeg interpretation. Springer Publishing Company, 376

[8] Klerk CC, S.V. Johnson MH: An eeg study on the somatotopic organisation of sensorimotor cortex activation during action execution and observation in infancy. Dev Cogn Neurosci, 376. https://doi.org/10.1016/j.dcn.2015.08.004

[9] Rojas GM, M.C.d.l.I.-V.M.C.J.G.M. Alvarez C: Study of resting-state functional connectivity networks using eeg electrodes position as seed. Front Neurosci (2018). https://doi.org/10.3389/fnins.2018.00235

[10] Jayant N Acharya 1, J.C.P.T.T.N.T. Abeer Hani: American clinical neurophysiology society guideline 2: Guidelines for standard electrode position nomenclature. ACM Trans. Embed. Comput. Syst. 4(33) (2016). https://doi.org/10.1097/WNP.0000000000000316

[11] Tiny Electronic Implants Monitor Brain Injury, Then Melt Away. https://news.illinois.edu/view/6367/312684

[12] Chi, Y.M., Jung, T.-P., Cauwenberghs, G.: Dry-contact and noncontact biopotential electrodes: Methodological review. IEEE Reviews in Biomedical Engineering 3, 106–119 (2010). https://doi.org/10.1109/RBME.2010.2084078

[13] Chi, Y.M., Cauwenberghs, G.: Wireless non-contact eeg/ecg electrodes for body sensor networks. In: 2010 International Conference on Body Sensor Networks, pp. 297–301 (2010). https://doi.org/10.1109/BSN.2010.52

[14] azhari, M.E., Toumanari, A., Latif, R.: Performance analysis of ieee 802.15.6 and ieee 802.15.4 for wireless body sensor networks. In: 2014 International Conference on Multimedia Computing and Systems (ICMCS), pp. 910–915 (2014). https://doi.org/10.1109/ICMCS.2014.6911180

[15] Manna, T., Misra, I.S.: Performance analysis of scheduled access mode of the ieee 802.15.6 mac protocol under non-ideal channel conditions. IEEE Transactions on Mobile Computing 19(4), 935–953 (2020). https://doi. org/10.1109/TMC.2019.2901852

[16] Huang, R., Nie, Z., Duan, C., Liu, Y., Jia, L., Wang, L.: Analysis and comparison of the ieee 802.15.4 and 802.15.6 wireless standards based on mac layer. Springer International Publishing, 7–16 (2015)

[17] Zhao, Z., Huang, S., Cai, J.: An analytical framework for ieee 802.15.6 based wireless body area networks with instantaneous delay constraints and shadowing interruptions. IEEE Transactions on Vehicular Technology PP(99), 1–1 (2018). https://doi.org/10.1109/TVT.2018.2789343

[18] Kahsay, L.Z., Paso, T., Iinatti, J.: Evaluation of ieee 802.15.6 mac user priorities with uwb phy for medical applications. In: 2013 7th International Symposium on Medical Information and Communication Technology (ISMICT), pp. 18–22 (2013). https://doi.org/10.1109/ISMICT.2013.6521691

[19] Sarkar, S., Misra, S., Bandyopadhyay, B., Chakraborty, C., Obaidat, M.S.: Performance analysis of ieee 802.15.6 mac protocol under non-ideal channel conditions and saturated traffic regime. IEEE Transactions on Computers 64(10), 2912–2925 (2015). https://doi.org/10.1109/TC.2015.2389806

[20] Yang, L., Li, C., Song, Y., Yuan, X., Lei, Y.: Performance evaluation of ieee 802.15.6 mac with user priorities for medical applications. In: Park, J.J.J.H., Pan, Y., Kim, C., Yang, Y. (eds.) Future Information Technology - II, pp. 233–240. Springer, Dordrecht (2015)

[21] Boulis, A., Tselishchev, Y., Libman, L., Smith, D., Hanlen, L.: Impact of wireless channel temporal variation on mac design for body area networks. ACM Trans. Embed. Comput. Syst. 11(S2), 51–15118 (2012). https://doi.org/10.1145/2331147.2331161

[22] El Azhari, M., El Moussaid, N., Latif, R.: Equalized energy consumption in wireless body area networks for a prolonged network lifetime. In: Wireless Communications and Mobile Computing, p. 9 (2017). https://doi.org/10.1155/2017/4157858. https://doi.org/10.1155/2017/4157858

[23] Mathapati, M., Kumaran, T.S., Prasad, K.H.S.: Framework with temporal attribute for secure data aggregation in sensor network. SN Applied Sciences 2(12), 2523–3971. https://doi.org/10.1007/s42452-020-03773-0

[24] Tselishchev, Y., Boulis, A.: Effects of sensor-to-sensor link modeling on body area network simulations. In: Proceedings of the 7th ACM Workshop on Performance Monitoring and Measurement of Heterogeneous Wireless and Wired Networks. PM2HW2N ‘12, pp. 191–198. ACM, New York, NY, USA (2012). https://doi.org/10.1145/2387191.2387217. http://doi.acm.org/10.1145/2387191.2387217

[25] Boulis, A., Tselishchev, Y.: Contention vs. polling: A study in body area networks mac design. In: Proceedings of the Fifth International Conference on Body Area Networks. BodyNets ‘10, pp. 98–104. ACM, New York, NY, USA (2010). https://doi.org/10.1145/2221924.2221944. http://doi.acm.org/10.1145/2221924.2221944

[26] Zhang, C., Ma, Y.: Ensemble machine learning: Methods and applications. Springer Publishing Company, Incorporated (2012)

[27] Alok Kumar, M.J.: Ensemble learning for ai developers (2020). https://doi.org/10.1007/978-1-4842-5940-5

[28] Jordan J. Bird, M.L.J.E.A. Diego Resende Faria: A deep evolutionary approach to bioinspired classifier optimisation for brain-machine interaction. https://doi.org/10.1155/2019/4316548

[29] Bird, J.J., Faria, D.R.: Mental emotional sentiment classification with an eeg-based brain-machine interface. (2018)

